# Morphological and thermoregulatory responses to urbanization in the European garden spider *Araneus diadematus*

**DOI:** 10.64898/2026.06.11.731659

**Authors:** Katrien De Wolf, Maxime Dahirel, Pieter Vantieghem, Bram Vanthournout, Mieke Soenens, Liliana D’Alba, Matthew Shawkey, Eva Vermeersch, Sylvia Lycke, Peter Vandenabeele, Dries Bonte

## Abstract

Urbanization creates novel environments that can drive phenotypic and behavioural responses, yet how multiple traits respond across spatial scales remains poorly understood. In particular, elevated ambient temperatures via the urban heat island effect may drive morphological and behavioural responses. We investigated body size, abdominal colouration, microhabitat use, behavioural thermoregulation and thermal offset relative to ambient air in the orb-weaving spider *Araneus diadematus* across rural-urban gradients in northern Belgium. Contrary to predictions from the temperature-size rule, body size increased with urbanization at large spatial scales, whereas size-corrected abdomen area—reflecting body condition and reproductive investment—declined with urbanization, with strongest support at local spatial scales. Abdominal colouration showed no response to urbanization despite evidence for both carotenoid-like pigments and melanin-associated structures. Nevertheless, body size and colouration covaried, with sites containing larger spiders tending to harbour darker individuals, whereas within sites larger individuals were slightly brighter than smaller conspecifics. Thermal responses showed little variation along the urbanization gradient. Retreats were consistently warmer than web hubs, and spiders maintained body temperatures above both their immediate microhabitat and ambient air. Only retreat-associated behavioural thermoregulation showed a weak decline with urbanization at local spatial scales. Our results reveal contrasting trait responses to urbanization across spatial scales and demonstrate that size-colour covariation can persist despite divergent responses of individual traits. These findings highlight the importance of considering multiple traits, their covariation and spatial scale to accurately understand and predict ecological responses of ectotherms to urban environments.

## 1. Introduction

Urbanization represents one of the most prominent forms of human-induced rapid environmental change, globally altering abiotic and biotic conditions (Parris, 2016; Sih et al., 2011). One of the most striking consequences is the urban heat island effect, whereby reduced vegetation cover, increased impervious surfaces and anthropogenic heat emissions elevate environmental temperatures relative to surrounding rural areas (Oke, 1973; Rizwan et al., 2008). Although the urban heat island intensity varies with city structure and regional climate (Manoli et al., 2019), even small urban centres often exhibit temperature increases of several degrees (Ward et al., 2016). Such thermal alterations can accelerate development and modify both maturation timing and size at maturity in ectotherms, with cascading consequences for life-history strategies, reproductive output and population dynamics (Alberti et al., 2017; Angilletta et al., 2002).

Morphological traits, including body size and colouration, often respond to ecological gradients associated with urban environments, such as thermal variation (Atkinson, 1994; Stuart-Fox et al., 2017). According to the temperature–size rule, elevated temperatures accelerate development relative to growth, resulting in smaller adult sizes in many ectotherms (Horne et al., 2015). Such size reductions parallel the inverse Bergmann’s rule observed in ectotherms across natural thermal gradients, where individuals tend to be smaller in warmer environments (Angilletta et al., 2004; Mousseau, 1997). However, if growth rates increase while phenology remains fixed, larger body sizes can occur despite higher temperatures, particularly when resource acquisition scales with activity (Frizot et al., 2025). Beyond thermal effects on growth, urban climates may also impose constraints on morphology, with desiccation risk favouring larger individuals while limits to heat dissipation favour small-bodied individuals (Nervo et al., 2021). Beyond climate, urban-induced habitat fragmentation may further influence phenotype-dependent dispersal (Cheptou et al., 2017; Fischer and Lindenmayer, 2007), although its importance for body size evolution is taxon-specific and may be less relevant for species where dispersal occurs primarily during juvenile stages, such as orb-web spiders (Bucher and Entling, 2011).

Beyond overall body size, condition-related traits may also respond to urban environmental changes. In spiders, abdominal size reflects energetic reserves that support reproduction and egg development and is therefore closely linked to individual condition and fitness (Jakob et al., 1996; Lowe et al., 2014; Moya-Laraño et al., 2008). Relative abdomen size may therefore respond differently to urbanization than overall body size, particularly if urban environments alter resource availability or energetic demands.

Colour variation across urbanization gradients may arise through multiple, non-exclusive processes. Colouration can influence thermal balance in ectotherms through thermal melanism, whereby darker individuals absorb more solar radiation and warm more rapidly, while lighter phenotypes may reduce overheating risk in warm environments (Clusella Trullas et al., 2007; Stuart-Fox et al., 2017). Urban colour shifts may also arise through non-thermal processes, including pollution-induced melanism or resource limitation that constrains pigment deposition, particularly for diet-derived pigments (Cook and Saccheri, 2013; Leveau, 2021; Salmón et al., 2021). Because colour is multifunctional, mediating crypsis, signalling and physiological protection (e.g. UV mitigation, antimicrobial defence and thermoregulation), it can be subject to multiple, and potentially opposing, selective pressures in urban environments (Oxford and Gillespie, 1998; Stuart-Fox et al., 2017).

Body size and colouration may jointly influence thermal balance under urbanization. Larger bodies have greater thermal inertia, meaning that identical increases in absorptivity (i.e. darker colouration) can produce higher peak and more sustained body temperatures. Conversely, combinations of smaller body size and lighter colouration may reduce overheating but increase desiccation risk. Selection may therefore act on trait (size–brightness) combinations rather than on individual traits alone (Clusella Trullas et al., 2007; Dahirel et al., 2025). Developmental factors such as temperature and nutrition may further link these traits by influencing both growth trajectories and pigment deposition.

In spiders, abdominal colouration is produced by both pigmentary compounds (e.g. ommochromes, bilins, eumelanin and carotenoids) and structural mechanisms such as guanine crystals and multilayer cuticular arrangements. These micro- to nanoscale photonic architectures determine how light is scattered, transmitted or reflected, thereby modifying brightness and hue and often amplifying or masking underlying pigments (Hsiung et al., 2017; Shawkey and D’Alba, 2017). Understanding the pigmentary basis is important because pigments differ markedly in origin and function: melanin is produced endogenously and contributes to UV protection and thermal absorption, whereas carotenoids are diet-derived and therefore linked to nutritional condition and resource availability, making them particularly sensitive to urban resource limitation and stressors (Grunst et al., 2020; Hsiung et al., 2015; Oxford and Gillespie, 1998). As a result, variation in colouration may reflect both environmental conditions and resource constraints in urban habitats. In orb-weavers, bright body patches can attract insect prey, while background matching may reduce detection by predators, indicating that colouration and colour patterns may influence ecological interactions (Peng et al., 2020; Robledo-Ospina et al., 2017; Ximenes et al., 2020, Messas et al., 2025). Yet, despite the ecological importance of pigmentary mechanisms and their likely sensitivity to urban environmental gradients, the biochemical basis of colour variation remains insufficiently characterised for many spiders (but see review Hsiung et al. (2019)), including those commonly found in cities.

In addition to morphological traits, behaviour provides a flexible and rapid mechanism for responding to environmental change (Sih et al., 2011; Tuomainen and Candolin, 2011; Wong and Candolin, 2015). In orb-weaving spiders, individuals may actively select thermally buffered sites for web construction or move between the exposed web hub and a nearby sheltered retreat. However, such behaviours may reduce prey interception and create trade-offs between occupying the optimal thermal niche and maintaining foraging efficiency (Huey et al., 2012; Taucare-Ríos et al., 2024). Behavioural adjustments such as posture changes, activity timing or web orientation, may further allow individuals to buffer temperature variation within microhabitats (Kearney et al., 2009; Woods et al., 2015).

To investigate how these traits and behaviours respond to urban environments, we focused on the orb-weaving spider *Araneus diadematus*, a widespread species occurring from semi-natural habitats to highly urban sites, including gardens and parks (Dahirel et al., 2019; Toft, 1976). This species exhibits pronounced abdominal colour variation, characterised by a conspicuous guanine-based dorsal cross set against a variable brownish-background ranging from pale yellow to near-black (Messas et al., 2025). The size and prominence of this white-coloured cross can vary substantially among individuals and populations (Seitz 1972; Blanke 1981), but it is unclear if this variation in colour pattern may be influenced by urbanization or other environmental changes. Previous work revealed that responses to urbanization may vary with spatial scale, particularly for morphology and web architecture, potentially as a consequence of urban-induced trophic changes (Dahirel et al., 2019). Whether colouration and thermoregulatory behaviour show similar responses to urbanization in this species remains unclear. Furthermore, little is known about covariation between body size and colouration.

Here, we investigated how morphology, colouration and thermoregulatory behaviour vary along the urbanization gradient. First, we analysed each trait independently to test for direct responses to urbanization. If the temperature–size rule applies under urban warming, then body size should decline with increasing urbanization. If colour-mediated thermoregulation occurs, then abdominal brightness should increase with urbanization, favouring lighter individuals in warmer environments. We further examined thermoregulatory behaviour by assessing microhabitat use and body temperatures relative to environmental conditions vary along the rural-urban gradient. Because body size and colouration may jointly influence thermal balance, we explicitly examined whether these traits covary in addition to testing their separate responses to urbanization. Finally, we characterized abdominal pigments to determine whether melanin or carotenoids underlie the observed colour variation, thereby clarifying the physiological basis of these colours and their potential ecological functions.

## 2. Materials and Methods

### 2.1. Study area and quantification of urbanization

Fieldwork was conducted in northern Belgium in 2022 across 54 local sites (200 x 200 m^2^), nested within 27 landscapes (3 x 3 km^2^). Each landscape contained two local sites selected to represent contrasting urbanization levels at the local scale (low: < 3% built-up area (BUA); high >15% BUA), allowing separation of local and landscape scale variation. This validated nested design ensured balanced sampling and maximized variation in urbanization at both local and landscape spatial scales (Merckx et al., 2018b). Urbanization was quantified as percentage BUA using the object-oriented Large-Scale Reference Database of Flanders (LRD, 2018), which delineates building footprints but excludes roads, public squares and parking lots. For each site, %BUA was calculated within twelve buffer radii ranging from 50 to 4000 m (50, 100, 200, 250, 400, 500, 800, 1000, 1600, 2000, 3200 and 4000 m) to capture variation across spatial extents (Figure 1).

**Figure 1:**
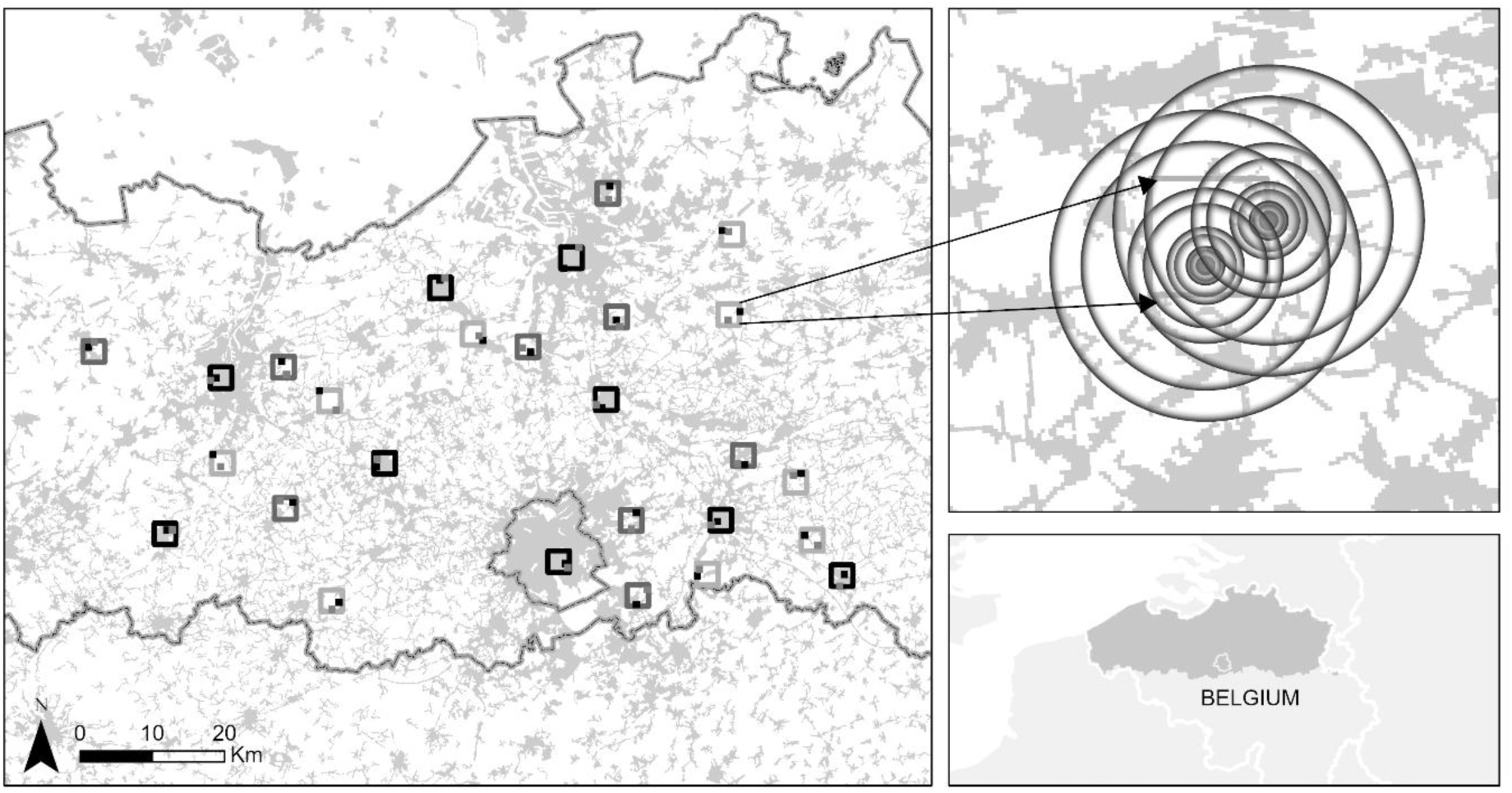
Map of the study area. Sampling was conducted across 54 local sites (200 × 200 m) nested within 27 landscapes (3 × 3 km) in northern Belgium. Squares indicate sampling sites distributed across the study region. Insets illustrate the nested structure of the design, with local sites paired within landscapes to capture variation at both local and landscape spatial scales. Urbanization was quantified as percentage built-up area (%BUA) within concentric buffers (50–4000 m) around each site (top inset). The lower panel shows the location of the study area within Belgium.

### 2.2. Microclimate and spider temperature measurements

At each site, temperature measurements were collected from up to five female spiders positioned in the web hub (i.e. centre of the web) and up to three individuals located in the retreat (i.e. hiding places adjacent to the web such as beneath vegetation or within curled leaves). Spider body surface temperature (T_spider_), ambient air temperature (T_ambient_) and microhabitat temperature in the direct vicinity of the web hub or in the retreat (T_webhub_ and T_retreat_) were measured using a T-type thermocouple (Omega T-type 36 gauge) connected to a handheld thermometer (Omega HH-25U), following methods described by Bladon et al. (2020). Measurements were repeated three times and mean values were used for analyses. Sampling occurred between 9 September and 6 October 2022. Because measurements were conducted across multiple days, weather conditions (e.g. cloud cover and solar radiation) were not identical among sites. To account for variation in absolute temperature among sampling days, thermal metrics were expressed as temperature differences relative to ambient air temperature.

We derived three thermal metrics: (1) Microclimate selection was calculated as the temperature difference between the microhabitat and ambient air (ΔT_webhub–ambient and_ ΔT_retreat–ambient_), indicating whether spiders occupy microhabitats that are warmer or cooler than ambient conditions. (2) Behavioural thermoregulation was calculated as the difference between spider body temperature and the temperature of its immediate microhabitat (ΔT_spider–webhub_ for individuals positioned in the web hub and ΔT_spider–retreat_ for retreat-positioned spiders), capturing the extent to which spiders maintained body temperatures that diverged from or matched their immediate surroundings. (3) Thermal offset was calculated as the difference between spider body temperature and ambient air temperature (ΔT_spider–ambient_), providing a standardized measure of deviation from ambient conditions and allowing comparisons of overall thermal exposure among sites.

### 2.3. Morphological measurements: body size and abdominal colouration

Collected spiders were photographed alive using a Nikon D3300 camera on a standardized background including a grey-scale card and a one-centimetre scale bar for colour and size calibration (Figure S2). Spider body size was approximated by measuring body length from the anterior cephalothorax to the posterior end of the opisthosoma (abdomen) using *ImageJ* (nearest 0.1 mm; Schneider et al., 2012). Abdomen area and dimensions of the guanine-based dorsal cross (width and length) were measured from the same images (Figure S2). Reflectance values from the grey-scale standards were obtained using an Avaspec-2048 spectrometer and converted to calibrated RGB reflectance using the R package *pavo* (Maia et al., 2013). Brightness was calculated as the arithmetic mean of the calibrated reflectance values, providing an estimate of lightness (achromatic intensity), with higher values corresponding to lighter or more reflective individuals (details in supporting information). Brightness was measured for three regions of the abdomen: (i) the whole abdomen, (ii) a standardized area of the white guanine cross, and (iii) a standardized area of the brown background.

To quantify variation in the prominence of the guanine-based cross, we calculated a cross prominence index as the ratio between the rectangular bounding box area of the cross (length × width) and the total abdomen area, providing a size-independent measure of the spatial extent of the cross pattern that allows standardized comparison among individuals.

Melanin detection was conducted using transmission electron microscopy (TEM). Spiders were preserved and fixed in a paraformaldehyde/glutaraldehyde solution. Following fixation, small pieces of the abdominal tegument were cut, post-fixed with 4% osmium tetroxide, dehydrated in an ascending series of ethanol and embedded in Epon. Sections were made on a Leica EM UC6 ultramicrotome at 100-150 nm and stained with 1% uranyl acetate and lead citrate. All observations were made using a JEOL JEM 1010 transmission electron microscope at 60 kV, equipped with a CCD side-mounted Veleta camera.

Pigment composition was also assessed using Raman spectroscopy (Bruker Optics Senterra dispersive Raman spectrometer equipped with an Olympus microscope and an XYZ motorized stage). Both 532 nm and 785 nm excitation lasers were evaluated and the 532 nm laser was selected for optimal signal. Spectra were acquired with a 20× objective, 10% laser power (1.58 mW), 10-s integration time and three accumulations. Spectral post-processing was conducted in *Thermo Grams/AI 9.0* (Thermo Fischer Scientific). β-carotene (PHR1239, Supelco®) was used as reference. Following preliminary tests, Raman spectra were acquired from abdominal tissue after cuticle removal, because colour variation was not retained in the isolated cuticles. Individuals spanning the full range of abdominal colour variation were selected for pigment analyses.

### 2.4. Statistical analysis

All analyses were conducted in R 4.5.2 (R Core Team, 2025).

To test how urbanization affects morphological traits (body length, abdomen area and spider colour) and thermal variables (microclimate selection, behavioural thermoregulation and thermal offset), we fitted linear mixed-effects models using *glmmTMB* (Brooks et al., 2017) with gaussian error distributions. Model assumptions were checked using *DHARMa* (Hartig, 2022) and response variables were log-transformed when necessary. All predictors were centered and scaled to improve interpretability and model convergence (Schielzeth, 2010), with scaling applied after log-transformation. Spider colouration was analysed using four response variables: cross prominence index, abdominal brightness, white cross brightness and brown background brightness. Because abdomen area was strongly correlated with body length (r_499_ = 0.931, p < 0.001), body condition was assessed by modelling log-transformed abdomen area as a function of %BUA while including log-transformed body length as a covariate, rather than using residuals (Moya-Laraño et al., 2008). The %BUA coefficient therefore represents the effect of urbanization on abdomen area after accounting for the expected allometric relationship between abdomen size and body length, with positive deviations indicating relatively larger abdomens for a given body length.

To evaluate spatial scale dependence, we fitted separate models for each of the twelve spatial radii (50-4000 m), with %BUA at the corresponding radius included as a continuous fixed effect and site identity as a random intercept. Sampling day did not covary with %BUA at any spatial scale (mean r52 ± SD = 0.066 ± 0.058), indicating that the temporal order of sampling was independent of the urbanization gradient. Because each site was sampled on a single day (except one site sampled twice, two days apart), sampling time was confounded with site identity and therefore not modelled explicitly. This multiscale approach follows Merckx et al. (2018b), and the spatial scale with the strongest support for each response variable was identified using AICc (Bartoń, 2025). For these models, we calculated marginal R^2^ (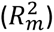) and conditional R^2^ (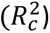), representing the variance explained by fixed effects and by both fixed and random effects, respectively (Nakagawa et al., 2017). Thermal responses were analysed using the thermal metrics defined above. Microclimate selection and behavioural thermoregulation were analysed separately for web hub and retreat-positioned individuals, whereas thermal offset was analysed across all individuals.

We further examined covariation between body size and colouration using a Bayesian multivariate mixed model implemented in the *brms* package (Bürkner, 2018) with the *cmdstanr* interface to the Stan language (Gabry et al., 2025; Stan Development Team, 2025). This approach allowed estimation of trait-specific responses to urbanization and the covariances between body size and colouration. As for the univariate analyses, separate models were fitted for each urbanization scale (50-4000 m radius) and the best-supported model was identified using leave-one-out cross validation (LOO) (Vehtari et al., 2017). In addition, a null model excluding urbanization was fitted to assess whether trait covariation was driven by shared responses to urbanization. Body size and abdominal brightness were modelled as above (scaled Gaussian response variables, log-transformed as needed, with urbanization (%BUA) as a fixed effect and site included as a random intercept). However, they were here modelled simultaneously, and the model also estimated correlations between traits at the site level (random-effect correlation) and at the within-site, individual level (residual correlation). Weakly informative Normal(0,1), Exponential(1) and LKJ(2) priors were used for fixed effects, standard deviations and correlations, respectively (McElreath, 2020). Models were fitted using Hamiltonian Monte Carlo by sampling four chains of 2000 iterations each, including 1000 warm-up iterations. Satisfactory convergence was confirmed following Vehtari et al. (2021) using R-hat statistics, effective sample sizes and visual inspection of trace plots.

## 3. Results

### 3.1. General

Across 54 sites, we collected thermal and morphological data for 455 female *A. diadematus* spiders (site mean ± SD: 8.43 ± 1.00; range per site: 4-10). Of these, 281 individuals were sampled in the web hub (5.20 ± 0.939; range: 2-7) and 174 in the retreat (3.22 ± 0.816; range: 1-6). Spider body surface temperatures ranged from 9.7° to 33.6°C, ambient air temperatures from 9.4°C to 27.4°C, web hub temperatures from 10.0° to 28.9°C and retreat temperatures from 9.7 to 29.8°C.

### 3.2. Morphological responses: body size and abdomen area

Urbanization at large spatial scales was strongly associated with spider size. Continuous models showed positive relationships between %BUA and body length from radii ≥ 400 m, with the 4000 m radius providing the best support (*p* < 0.001; Table 1; full details in Table S1). A 25% increase in %BUA within the surrounding 4000 m radius corresponded to an increase of 2.72 mm in body length and 28.04 mm^2^ in abdomen area, corresponding to increases of 26.8% ± 4.7% and 82.6% ± 17.5%, respectively (Figure 2). This allometric change indicates a slightly steeper increase in abdomen area relative to body length. However, the difference remains within the geometric expectations (a 27% increase in linear dimension approximates a 61% increase in area) (Table 1). At smaller spatial scales (≤ 200m), body size was either unaffected or negatively associated with urbanization. After accounting for body length, size-corrected abdomen area decreased significantly with increasing %BUA across all spatial scales (p < 0.05 at most scales; marginal negative effect at 4000 m, *p* = 0.086; Table S1). Thus, spider body length and size-corrected abdomen area showed contrasting responses to urbanization across spatial scales.

**Figure 2:**
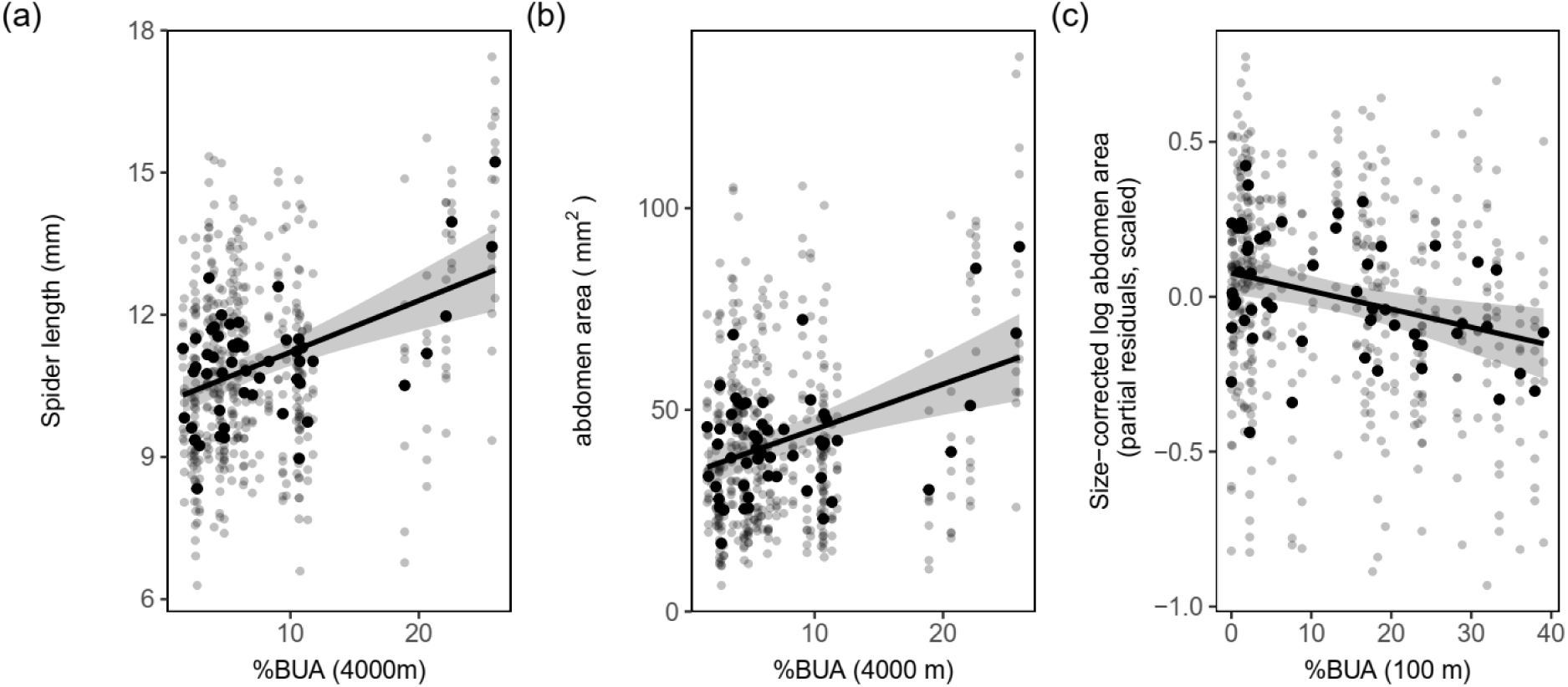
Relationship between urbanization and spider morphology. Spider body length (A) and abdomen area (B) increased with percentage built-up area (%BUA) at 4000 m. Size-corrected abdomen area (log-transformed partial residuals) (C) decreased with %BUA at 100 m. Grey points show individual measurements (n=455) and black points show site means. Black lines show fitted linear mixed-effects models and shaded regions represent 95% confidence intervals.

**Table 1:** Effects of urbanization (%BUA) on morphological traits and thermal metrics. The estimate for the percentage built-up area (%BUA) at the spatial scale (radius in metres) of the best-supported model is reported with its standard error, p-value, marginal and conditional R^2^. Models with significant effects of %BUA are shown in bold.

| Response variables | %BUA estimate | Std. error | z value | P value | $R_m^2 ; R_c^2$ | Scale (range) (m radius) |
| --- | --- | --- | --- | --- | --- | --- |
| **Morphological variables** |  |  |  |  |  |  |
| Body length | **0.339** | **0.072** | **4.7** | **<0.001** | **0.115 ; 0.311** | **4000 (400-4000)** |
| Abdomen area | **0.309** | **0.079** | **3.898** | **<0.001** | **0.096 ; 0.354** | **4000 (800-4000)** |
| Abdomen area (log-transformed; body length as covariate)<sup>a</sup> | **-0.071** | **0.025** | **-2.88** | **0.004** | **0.895 ; 0.918** | **100 (50-3200)** |
| Cross prominence index | -0.084 | 0.052 | -1.623 | 0.105 | 0.007 ; 0.028 | 4000 |
| Abdominal brightness (log-transformed) | -0.026 | 0.088 | -0.299 | 0.765 | 0.001 ; 0.328 | 200 |
| White cross brightness (log-transformed) | -0.12 | 0.071 | -1.7 | 0.092 | 0.015 ; 0.19 | 200 |
| Brown background brightness (log-transformed) | 0.089 | 0.078 | 1.142 | 0.254 | 0.008 ; 0.243 | 1000 |
| **Thermal variables** |  |  |  |  |  |  |
| $\Delta T_{\text{webhub-ambient}}$ (microclimate selection at web hub) | 0.052 | 0.055 | 0.948 | 0.343 | 0.003 ; 0.053 | 100 |
| $\Delta T_{\text{retreat-ambient}}$ (microclimate selection at retreat) | 0.086 | 0.058 | 1.486 | 0.137 | 0.007 ; 0.079 | 200 |
| $\Delta T_{\text{spider-webhub}}$ (behavioural thermoregulation for spiders positioned in the web hub ) | 0.110 | 0.09 | 1.214 | 0.225 | 0.012 ; 0.316 | 1000 |
| $\Delta T_{\text{spider-retreat}}$ (behavioural thermoregulation for spiders positioned in the retreat) | **-0.206** | **0.09** | **-2.284** | **0.022** | **0.043 ; 0.251** | **100 (100-400)** |
| $\Delta T_{\text{spider-ambient}}$ (Thermal offset)<sup>b</sup> | -0.043 | 0.075 | -0.567 | 0.571 | 0.017 ; 0.230 | 100 |

a: body size–corrected abdomen area was modelled as log-transformed abdomen area and log-transformed body length was included as a covariate. $R^2$ values reflect total variance in abdomen area explained by the model, including by body length.
b: position spider in the web (web hub or retreat) included as a covariate ( $R^2$ values again reflect variance explained by all predictors).

### 3.3. Morphological responses: abdominal colouration

The cross prominence index did not vary with %BUA at any spatial scale (Table S1). Likewise, abdominal brightness of the whole abdomen, white cross and brown background showed no association with urbanization (Table S1; Figure S5). Individuals nevertheless spanned a broad range of brightness values, from very pale to very dark phenotypes (Table S3). Raman spectroscopy of abdominal samples revealed consistent spectral peaks at approximately 1007, 1155 and 1513 cm⁻¹ across individuals, matching the β-carotene reference standard (Figure 3). These peaks were present across the full range of abdominal colour phenotypes, from pale yellow to dark brown. Transmission electron microscopy further revealed a distinct layer of highly electron-dense granules in the epidermal region of the abdominal cuticle, consistent with melanosomes (Figure 4). Together, these results indicate contributions of both carotenoid-like pigments and melanin-associated structures.

**Figure 3.**
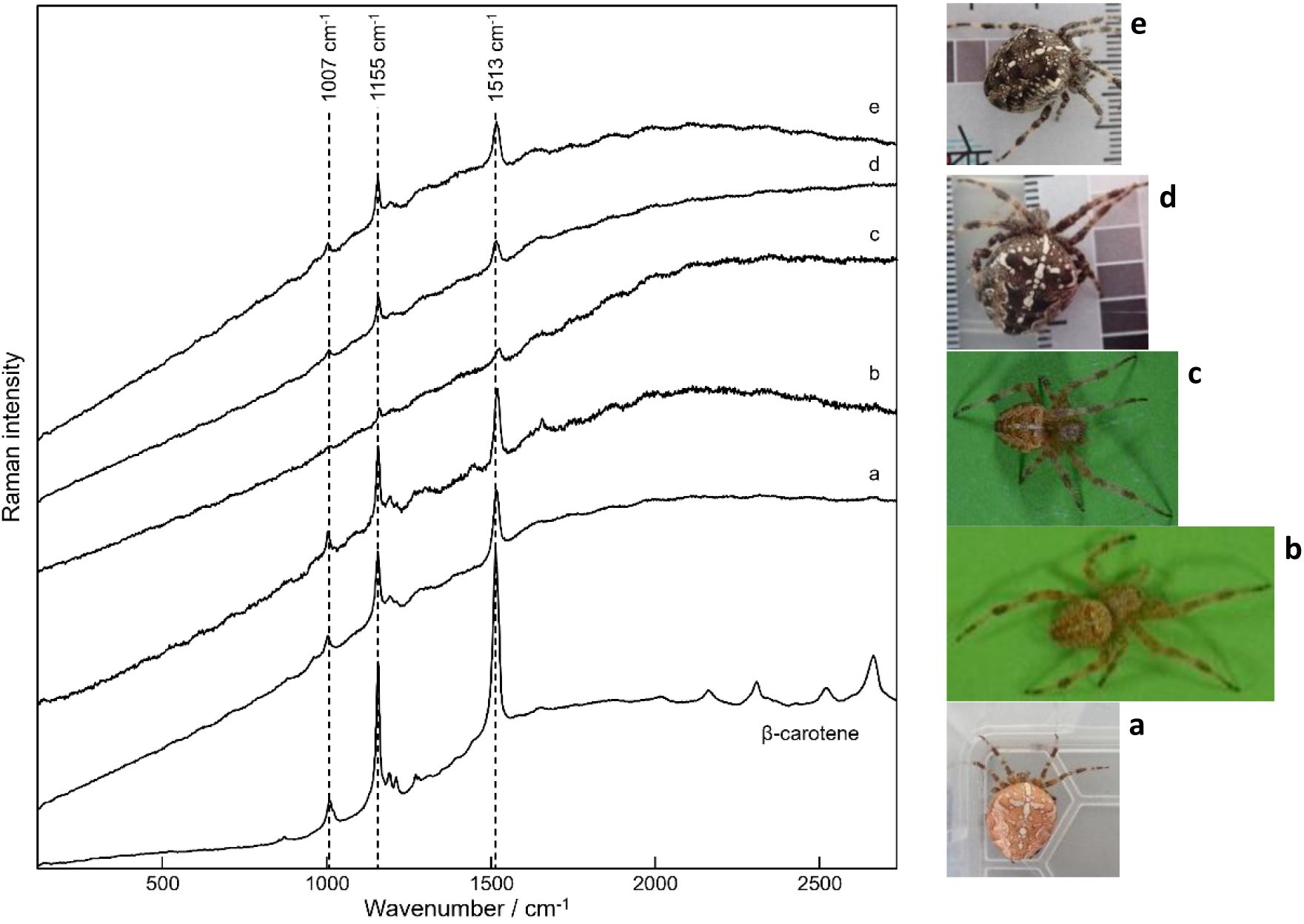
Raman spectra of abdominal colour variants. Spectra from five *A. diadematus* individuals spanning the observed colour range: (a) pale orange, (b) pale yellow, (c) pale brown, (d) dark brown and (e) black. All spectra show peaks at approximately 1007, 1155 and 1513 cm⁻¹, matching the β-carotene reference standard (PHR1239, Supelco®).

**Figure 4.**
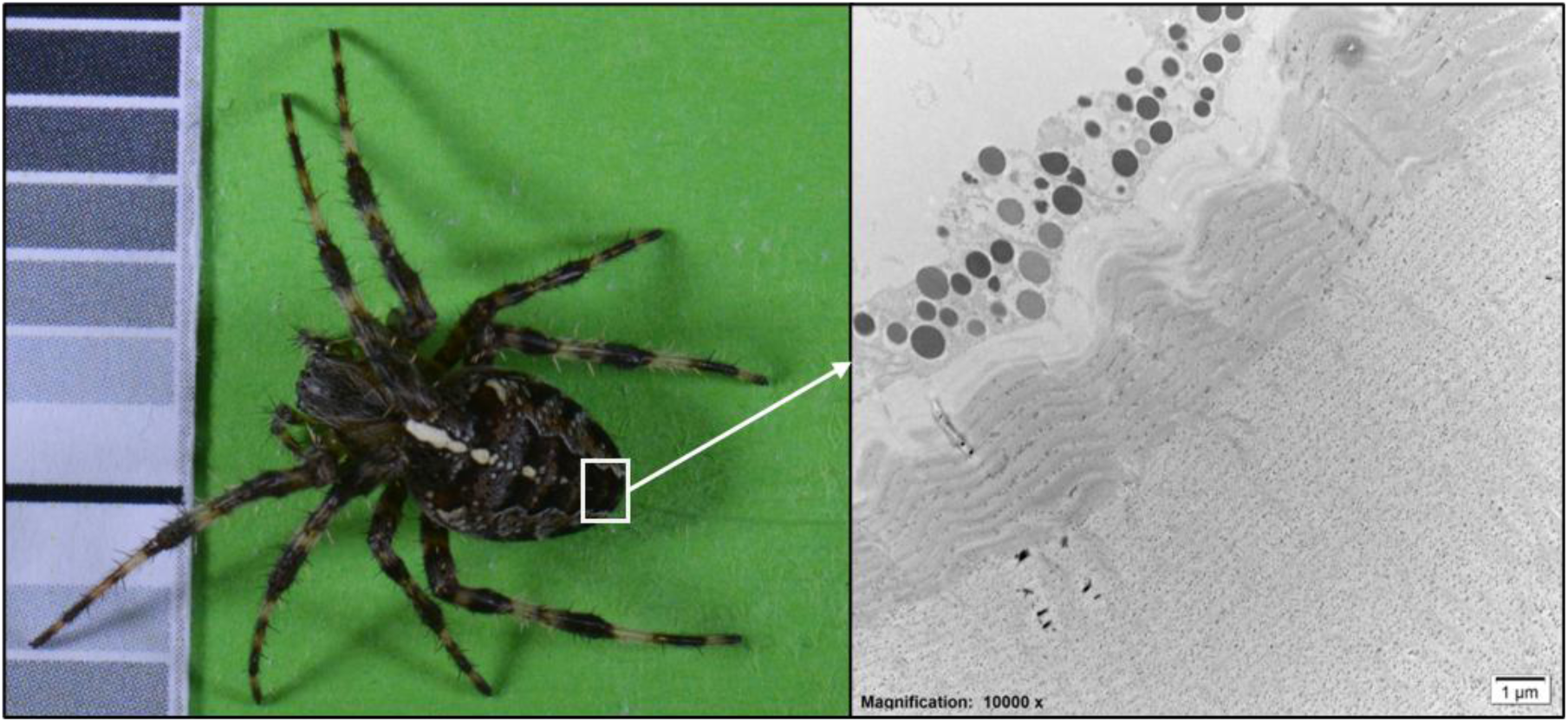
Transmission electron micrograph of abdominal melanosomes. Dorsal view of a spider, white square indicates the sampled abdominal region. TEM image showing a subcuticular layer of highly electron-dense granules, melanosomes. TEM magnification: 10,000x; scale bar: 1µm.

### 3.4. Morphological covariation: body size and colour

Body size and abdominal colouration showed covariation (Table 2). Consistent with the univariate analyses, urbanization had no effect on abdominal brightness, whereas body size increased with urbanization (β = 0.34 [95% credible interval: 0.19, 0.49]). At the site level, body length and abdominal brightness were negatively correlated after accounting for the fixed effect of urbanization (r[sites] = - 0.51 [-0.78, -0.19]), indicating that sites with larger spiders tended to host darker individuals. Within sites, however, the residual correlation between traits was positive (r[spiders] = 0.24 [0.15, 0.33]), suggesting that larger individuals were slightly brighter than smaller conspecifics. Variation in both traits was primarily driven by within-site (residual) variability, with more limited variation among sites (Table 2). Comparison of models fitted across urbanization scales using leave-one-out cross-validation identified the 4000 m radius model as the best-supported model, consistent with the scale-dependent body-size response observed in the univariate analyses (Table S2). The covariance structure between body size and abdominal brightness remained highly consistent across urbanization scales and was qualitatively similar to a model excluding urbanization (r[sites] = -0.43 [-0.68, -0.13] and r[spiders] = 0.22 [0.13, 0.31]), indicating that the observed trait associations were not driven solely by shared responses to urbanization (Table S2).

**Table 2.** Posterior summary of the Bayesian multivariate mixed model analysing the effect of urbanization (%BUA at 4000 m) on spider body size (body length) and spider colour (abdominal brightness). Both traits were modelled jointly to estimate their responses to urbanization and their covariation. Values represent posterior means with 95% credible intervals. The correlation between traits at the site level and at the individual (residual) level are also given.

|  | Body length (scaled) | Abdominal brightness (scaled) |
| --- | --- | --- |
| Fixed effects |  |  |
| Intercept | -0 (-0.15 – 0.14) | 0.01 (-0.16 – 0.18) |
| %BUA | 0.34 (0.19 – 0.49) | -0.01 (-0.18 – 0.16) |
| Random effects |  |  |
| Site-level SD | 0.46 (0.35 – 0.60) | 0.58 (0.45 – 0.74) |
| Distributional parameters |  |  |
| Residual SD | 0.83 (0.78 – 0.89) | 0.79 (0.73 – 0.85) |
| Bayesian R<sup>2</sup> |  |  |
| Full model predictions | 0.31 (0.24 – 0.37) | 0.31 (0.24 – 0.38) |
| Fixed effects only | 0.12 (0.04 – 0.21) | 0.01 (0 – 0.04) |
| Trait covariance |  |  |
| Individual-level (residual) correlation | 0.25 (0.15 – 0.35) |  |
| Site-level correlation | -0.51 (-0.77 – -0.18) |  |

### 3.5. Thermal responses

Urbanization (%BUA at 50–4000 m) did not explain variation in microclimate selection. Web hubs closely matched ambient air temperature (ΔT_webhub–ambient_ ≈ 0; Figure S3A), whereas retreats were consistently warmer (ΔT_retreat–ambient_ > 0; Figure S3B). Across sites, retreats were on average 0.46°C ± 0.06°C warmer than web hubs (p = 0.004), a difference unrelated to urbanization.

Spiders positioned at web hubs maintained body temperatures above the surrounding web microclimate, with an average thermoregulatory elevation of 2.93°C (intercept-only model, 95% CI: 2.69-3.18, *p* < 0.001). However, behavioural thermoregulation showed no relationship with urbanization at any spatial scale (all *p* > 0.1; Table 1; Table S1). Retreat-positioned spiders likewise maintained body temperatures above retreat microclimates, although this difference decreased with increasing %BUA at small spatial scales (100-250 m), with strongest support at 100 m (*p* = 0.022; Table 1; Figure S4), although the proportion of variance explained was low 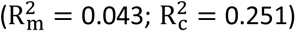. On average, spiders maintained body temperatures above ambient air temperatures. This thermal offset showed no relationship with urbanization at any spatial scale (all p > 0.05; Table S1). However, spiders positioned in the web hub had higher thermal offsets than those in retreats, with body temperatures 0.41°C higher relative to ambient air (95% CI: 0.14-0.67, *p* = 0.003).

## 4. Discussion

Urbanization was associated with contrasting responses across phenotypic traits and spatial scales in *A. diadematus*. Body size (total body length and abdomen area) increased with urbanization at large spatial scales, whereas at smaller spatial scales body size effects were weak or negative. Size-corrected abdomen area declined along the urbanization gradient, indicating reduced energetic reserves or delayed maturation. In contrast, abdominal colouration, microhabitat selection and behavioural thermoregulation showed little variation along the rural-urban gradient. Together, these results suggest that urbanization does not induce uniform phenotypic shifts.

Body length and abdomen area exhibited the strongest responses to urbanization, with individuals in more urbanized landscapes being larger. This result contrasts with expectations from the temperature–size rule (Horne et al., 2017, 2015), and with previous findings from the same region reporting reduced body size at highly urbanized local sites (Dahirel et al., 2019). These differences may arise from variation in spatial scale and trait definitions between studies, as Dahirel et al. (2019) modelled simultaneous responses to urbanization at two spatial scales (local and landscape scale) and used cephalothorax width to measure body size, whereas we examined multiple spatial extents to retain the single best-supported one, and used body length. More broadly, our findings support the view that body size responses to thermal environments are context-dependent and often deviate from ecogeographical expectations, including the inverse Bergmann’s rule commonly observed in ectotherms (Goldenberg et al., 2022). Positive size shifts have also been reported in other urban spiders, such as *Nephila plumipes*, and in arthropods more generally, where larger body size may enhance dispersal, colonization success or competitive ability (Lowe et al., 2014; Merckx et al., 2018a, 2018b). However, because *A. diadematus* disperses during juvenile stages and adults show strong site fidelity (Bonte et al., 2023), fragmentation-driven filtering is unlikely to explain the size patterns observed here. Instead, responses to warming at landscape scales may override local constraints, potentially via accelerated growth or extended activity periods that enhance resource assimilation. Although the urban heat island effect is strongest at local spatial scales (200-500 m), typically increasing near-surface air temperatures by ∼1-2°C on average, it extends across kilometres, including the 4 km landscape considered here (Caluwaerts et al., 2020; Merckx et al., 2018b; Ward et al., 2016). Such variation across spatial scales may contribute to faster growth and larger size at landscape scales, while local thermal extremes and resource limitation may impose opposing pressures.

In contrast to body size, relative abdomen area decreased after accounting for body length, suggesting reduced body condition in more urbanized environments. Because abdomen area is closely linked to body condition and fecundity in spiders (Jakob et al., 1996; Lowe et al., 2014; Moya-Laraño et al., 2008), this result suggests that local urban environments may constrain resource acquisition or increase energetic demands. Reduced prey availability in urban environments at the local scale likely contributes to this pattern, as decreases in insect biomass and shifts in prey composition have been reported at the same locations in a sampling campaign in 2021 (De Wolf et al., 2025). Temperature extremes may also contribute by altering prey activity and availability, or by affecting spider performance and energetic demands (Kingsolver et al., 2013). However, we cannot formally disentangle thermal and trophic effects because temperature was measured only on the sampling day and therefore was not integrated into the trait models. Given that sampling occurred during the reproductive period, smaller abdomens probably reflect lower egg load or delayed maturation, as previously observed in urban populations of *A. diadematus* (Dahirel et al., 2019).

Abdominal colouration showed no detectable response to urbanization, despite clear evidence that colour is produced by pigments that differ in their sensitivity to nutritional condition (e.g. β-carotene vs melanin). Raman spectroscopy revealed β-carotene signatures and transmission electron microscopy confirmed melanosomes, yet brightness and cross prominence remained stable across the rural-urban gradient. This stability likely reflects multifunctional and potentially antagonistic selection on colour traits, such as prey attraction versus crypsis, which may constrain directional shifts when thermoregulatory benefits are modest relative to other ecological functions (e.g., signalling or camouflage). Urban environments may further reinforce this pattern by generating heterogeneous visual backgrounds and predator communities, promoting stabilizing rather than directional selection. Consistent with this interpretation, colour responses to urbanization are often context-dependent across taxa and shaped by multiple, non-exclusive drivers. Urban colour shifts have been linked to pollution-driven melanism and/or background matching rather than temperature alone (Keinath et al., 2020). More broadly, large-scale analyses show that the effectiveness of conspicuous versus cryptic colour strategies depends on predator communities, light environments and prey composition, rather than following a single directional trend (Medina et al., 2025). Although the thermal melanism hypothesis predicts darker individuals in cooler environments due to faster heat gain, empirical support is mixed and frequently confounded by these alternative functions of colouration, including camouflage and protection against environmental stressors (Clusella-Trullas et al., 2008). Recent work suggests that camouflage in *A. diadematus* is achieved through behavioural substrate choice and background matching (Messas et al., 2025), potentially reducing selection for directional changes in colouration along rural-urban gradients.

Body size and abdominal colouration were not independent but showed contrasting patterns of covariation within and among sites. At the individual level, larger spiders tended to be slightly brighter, whereas at the site level populations with larger spiders were, on average, darker. These opposing patterns suggest that the relationship between size and colouration may reflect different processes operating within and among populations. Such scale-dependent covariance is expected when different processes shape trait associations within and among populations (Peiman and Robinson, 2017). Covariation among individuals may arise through shared developmental pathways or differences in resource acquisition, whereas among-population patterns may reflect environmental variation, local adaptation, plastic responses or pleiotropy. Individual-level covariation may therefore be linked to condition-dependent pigmentation or allometric relationships, whereas site-level patterns may reflect environmental influences on both traits. The negative site-level association further suggests that environmental drivers generating larger body sizes may also influence colouration, either directly or indirectly. For example, variation in prey availability, habitat structure, microclimate, or predator communities could favour different size–colour combinations among populations. In cryptic prey, body size can modify the detectability and relative performance of colour variants, creating correlational selection on trait combinations rather than on individual traits alone (Karpestam et al., 2014)

The site-level covariance also has implications for responses to urbanization. If body size and colouration were strongly linked through genetic covariance, evolutionary responses would be expected to occur preferentially along this covariance axis, potentially constraining independent trait evolution (Peiman and Robinson, 2017). However, body size increased with urbanization whereas abdominal brightness showed no detectable urban response, suggesting that responses of the two traits are at least partially decoupled. This interpretation is further supported by the near-identical covariance structure observed in models excluding urbanization, indicating that the opposing within-and among-site correlations are largely independent of urbanization itself. While we cannot distinguish between plastic and genetic mechanisms, this pattern suggests that the size–colour association is unlikely to strongly constrain phenotypic responses. Although the underlying mechanisms remain unclear, these results indicate that body size and colouration do not vary independently and that analyses focusing on single traits may overlook important aspects of phenotypic responses to urban environments.

Thermal variables showed limited variation along the rural-urban gradient. Microhabitat temperatures differed consistently between web hubs and retreats, but these differences were not associated with urbanization. Similarly, spiders maintained body temperatures above both their immediate microhabitat and ambient air, but the magnitude of these differences showed little variation along the urbanization gradient. This suggests that web placement may be influenced more strongly by structural constraints and foraging requirements rather than by thermal conditions. However, we did not test whether spiders adjust their use of retreats and web hubs to modulate thermal exposure. Such behaviour could buffer thermal differences among environments, resulting in similar patterns across the urbanization gradient, but may also reduce prey interception, generating trade-offs between thermal safety and foraging efficiency (Huey et al., 2012; Taucare-Ríos et al., 2024). Because *A. diadematus* is active during both day and night, and the urban heat island effects are often strongest after sunset, nocturnal dynamics may additionally contribute to urban-rural differences that were not captured by our daytime measurements. Across all sites, thermal offsets remained positive, indicating that spiders were consistently warmer than ambient air regardless of urbanization.

In summary, urbanization was associated with increased body size at large spatial scales, with spiders in highly urbanized landscapes being larger, although relative abdomen area declined after accounting for body size. However, this enlargement did not correspond to meaningful changes in thermal exposure, and colour traits remained stable along the rural-urban gradient. Importantly, body size and colouration remained covaried despite their contrasting responses to urbanization, indicating that trait relationships may persist even when individual traits respond differently to environmental change. Together, these findings highlight the importance of considering multiple traits, their covariation and spatial scale when predicting ectotherm responses to urban environments.

## Acknowledgements

We thank all cooperating citizens for granting access to their private gardens. We thank the Zoo of Antwerp for providing a sampling location. We thank Thomas Roels, Lukas De Jaegher and Laura Buonafede for their assistance with the field data collection. We also thank Bor-Kai Hsiung for advice on conducting Raman spectroscopy.

## Funding information

This study has received funding from the Research Foundation – Flanders (FWO grant G080221N to D.B. and GOG2217N to M. Shawkey), the Air Force Office of Scientific Research (AFOSR FA9550-23-1-0822 and FA9550-18-1-0447 to M. Shawkey) and the European Office of Aerospace Research and Development (EOARD FA8655-23-2-7041 to M. Shawkey)

## Author contributions

K.D.W.: Conceptualization, Methodology, Investigation, Data curation, Formal analysis, Visualization, Project administration, Writing – original draft. M.D.: Methodology, Validation, Writing – Review and Editing. P. Vantieghem: Conceptualization, Investigation. B.V.: Conceptualization, Writing – Review and Editing. M. Soenens: Investigation, Writing – review and editing. L.D.: Methodology, Writing – Review and Editing. M. Shawkey: Conceptualization, Funding acquisition, Writing – review and editing. E V.: Methodology, Investigation, Visualization, Writing – review and editing. S. L.: Methodology, Resources. Writing – review and editing. P. Vandenabeele.: Methodology, Resources, Writing – review and editing. D. B.: Conceptualization, Methodology, Supervision, Funding acquisition, Writing – review and editing.

## Conflict of interest statement

The authors declare no conflict of interest.

## Data availability statement

Data and code supporting the results of this study are available on OSF: https://osf.io/ez3b4/overview?view_only=5d3a51fdf55c4939b59f8fd2a95e2ab8

## Notes

### Competing Interest Statement

The authors have declared no competing interest.

