## Supplementary Material for "Morphological and thermoregulatory responses to urbanization in the European garden spider *Araneus diadematus*"

### Supporting Materials and Methods

**Colour quantification**

Reflectance values of the grey-scale calibration card were determined using an AvaSpec-2048 spectrometer and a dual light-source set-up (AvaLight-DH-S deuterium halogen and AvaLight-HAL-S-MINI light source), a bifurcated probe and an integrating sphere (AvaSphere-50-REFL). These measurements were used to convert the raw RGB values from photographs into calibrated reflectance values using the R package *pavo* (Maia et al., 2019). Each spider was photographed on a standardized background containing the grey-scale card and a 1-cm scale bar. For each image, average RGB values were extracted for each patch of the grey-scale (i.e. one white rectangle, one black rectangle and seven shades of grey rectangles) and for the selected abdominal regions using ImageJ (Schneider et al., 2012). Calibration curves were fitted separately for each colour channel (R,G and B), linking RGB intensity values to measured reflectance following the method of Johnsen (2016). Calibration performance was high for all colour channels. The mean correlation between observed and predicted reflectance values was 0.998 (range: 0.993 – 1). Brightness was calculated as the arithmetic mean of the reflectance values across the three colour channels and provides an estimate of the achromatic intensity across the visible spectrum of the spider abdomen, with higher values corresponding to brighter individuals (Stevens et al., 2014).

**Raman spectrometry**

Micro-Raman spectroscopy was performed using a Bruker Optics *Senterra* dispersive Raman spectrometer equipped with an Olympus microscope and an XYZ motorized stage for precise laser focusing. The spectrometer has two laser wavelengths: a 532 nm Nd:YAG laser and a 785 nm diode laser. Neutral density filters are used to control the laser power ranging between 0.03 (1%) and 14.34 mW (100%) or 0.11 (1%) and 45.2 mW (100%), for respectively the 532 and 785 nm laser. The detection is achieved by using a thermoelectrically cooled charge-coupled-device (CCD) detector, operating at a temperature of -65 °C. Spectra can be obtained with a spectral resolution of 3 to 5 $cm^{-1}$

in the range between 50 and 3700 $cm^{-1}$ and between 80 and 3500 $cm^{-1}$ for respectively the 532 and 785 nm laser. The microscope turret is equipped with objectives ranging from 5x (numerical aperture (NA): 0.1), 20x (NA: 0.4), 50x (NA: 0.75) to 100x (NA: 0.9) magnification. Respectively, the following spot sizes can be obtained: 50 µm, 10 µm, 4 µm and 2 µm (visual inspection, 785 nm laser). The system is controlled by OPUS software (Bruker). Both laser wavelengths were initially tested to determine optimal signal quality. The 532 nm laser was selected for all subsequent measurements due to superior signal-to-noise ratios. The laser power was set to 10% (1.58 mW) as to avoid degradation or transformation of the sample during acquisition. The spectra were acquired using the 20x objective with an integration time of 10 seconds and three accumulations per measurement. Post-spectral processing was done using Thermo Grams/AI 9.0 suite software (Thermo Fischer Scientific). β-carotene (PHR1239, Supelco®) was used as a reference standard. To ensure pigment signals were not obscured by structural components of the cuticle and as the isolated cuticles no longer retained visible colour variation, Raman measurements were taken of the abdominal tissue following cuticle removal.

### Supporting Figures

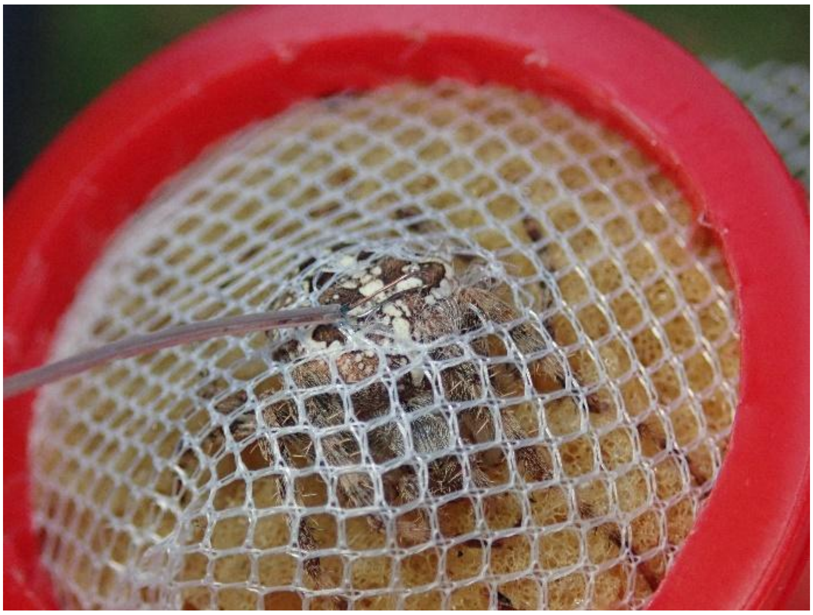

**Figure S1: Measurement of spider body surface temperature.**
Illustration of the procedure used to measure spider body surface temperature ($T_{spider}$) in the field. A fine-gauge T-type thermocouple (Omega® Type T, 36-gauge) was gently positioned against the dorsal surface of the abdomen.

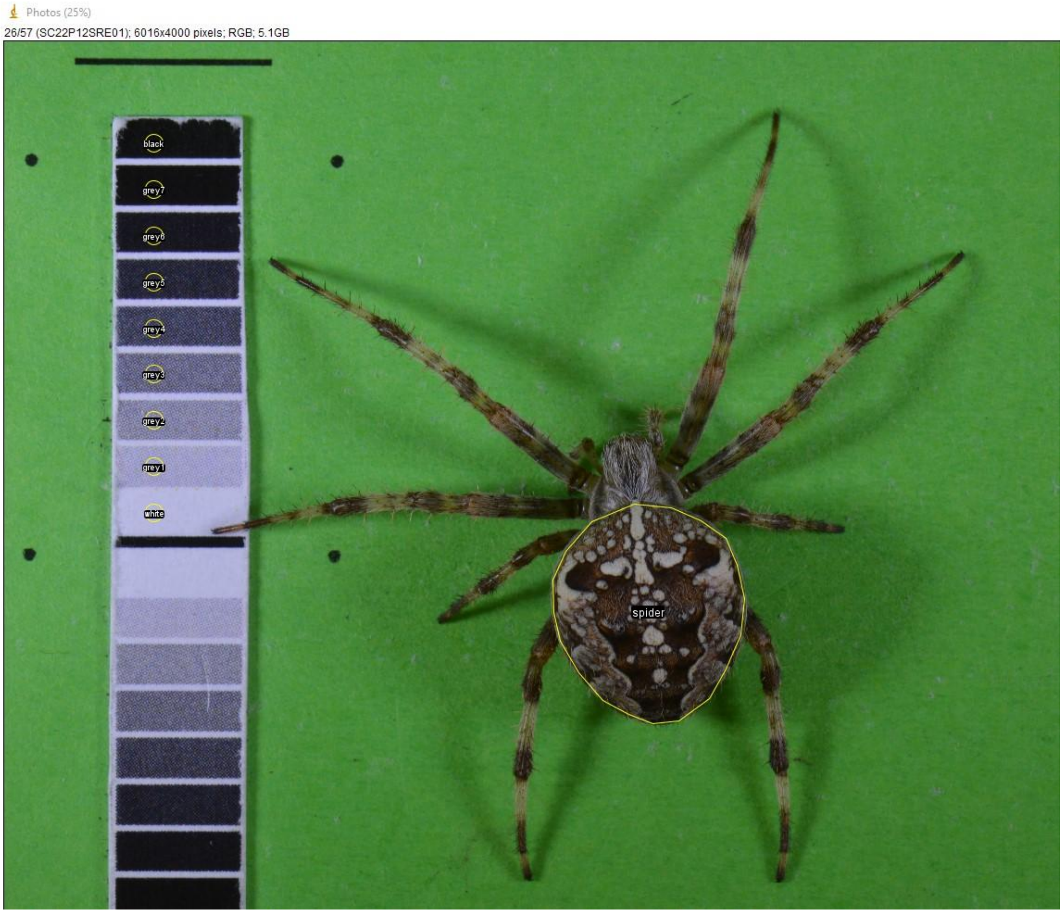


**Figure S2: Calibration setup.**

Example photograph of a live *A. diadematus* on a standardized background with a grey-scale calibration card and a 1 cm scale bar. The grey-scale card contained one white, one black and seven shades of grey rectangles. The image illustrates the selected whole-abdomen region and the selected grey-card regions for colour calibration.

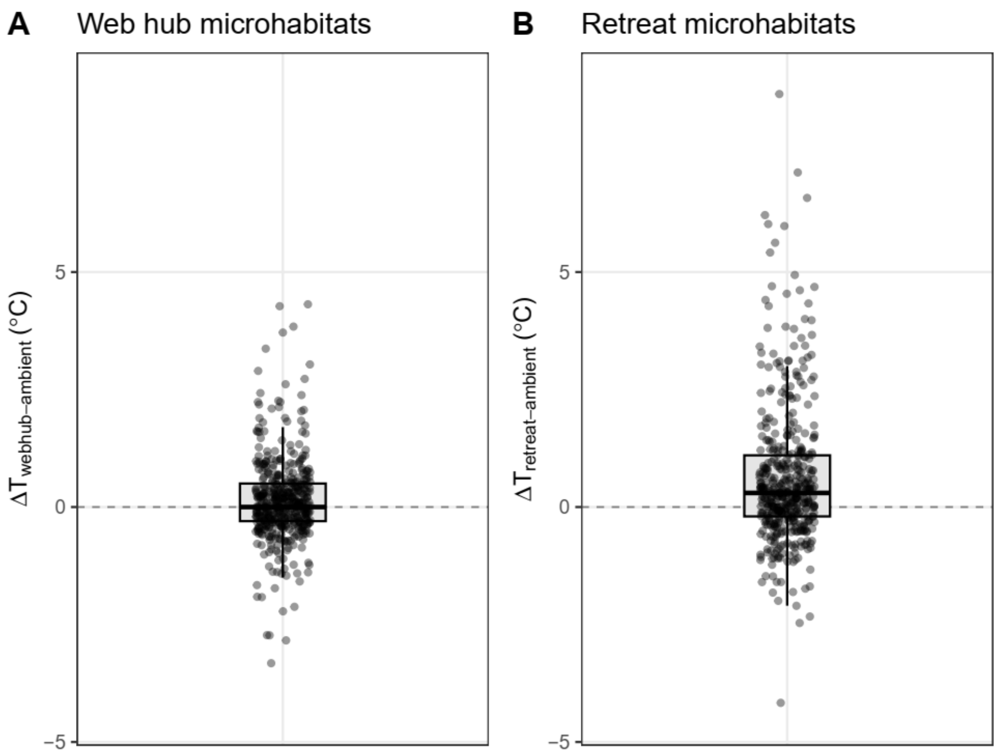


**Figure S3. Microhabitat temperature differences.**

(A) Temperature difference between the web hub and ambient air ($\Delta T_{webhub-ambient}$) across all sites.

(B) Temperature difference between the retreat and ambient air ($\Delta T_{retreat-ambient}$) across all sites.

Points represent individual spiders.

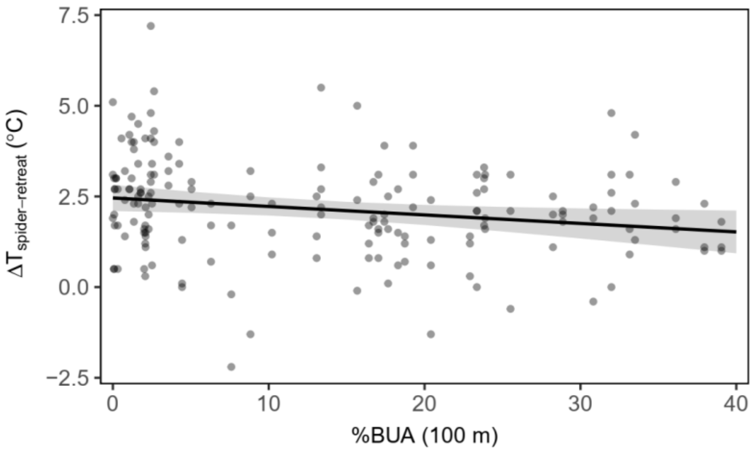


**Figure S4: Relationship between urbanization and behavioural thermoregulation of retreat positioned spiders.**

Temperature difference between spiders body surface temperature and retreat temperature ($\Delta T_{spider-retreat}$) in relation to percentage built-up area (%BUA) at 100 m radius. Points show individual measurements (n = 174). Black lines show fitted linear mixed-effects models and shaded regions represent 95% confidence intervals.

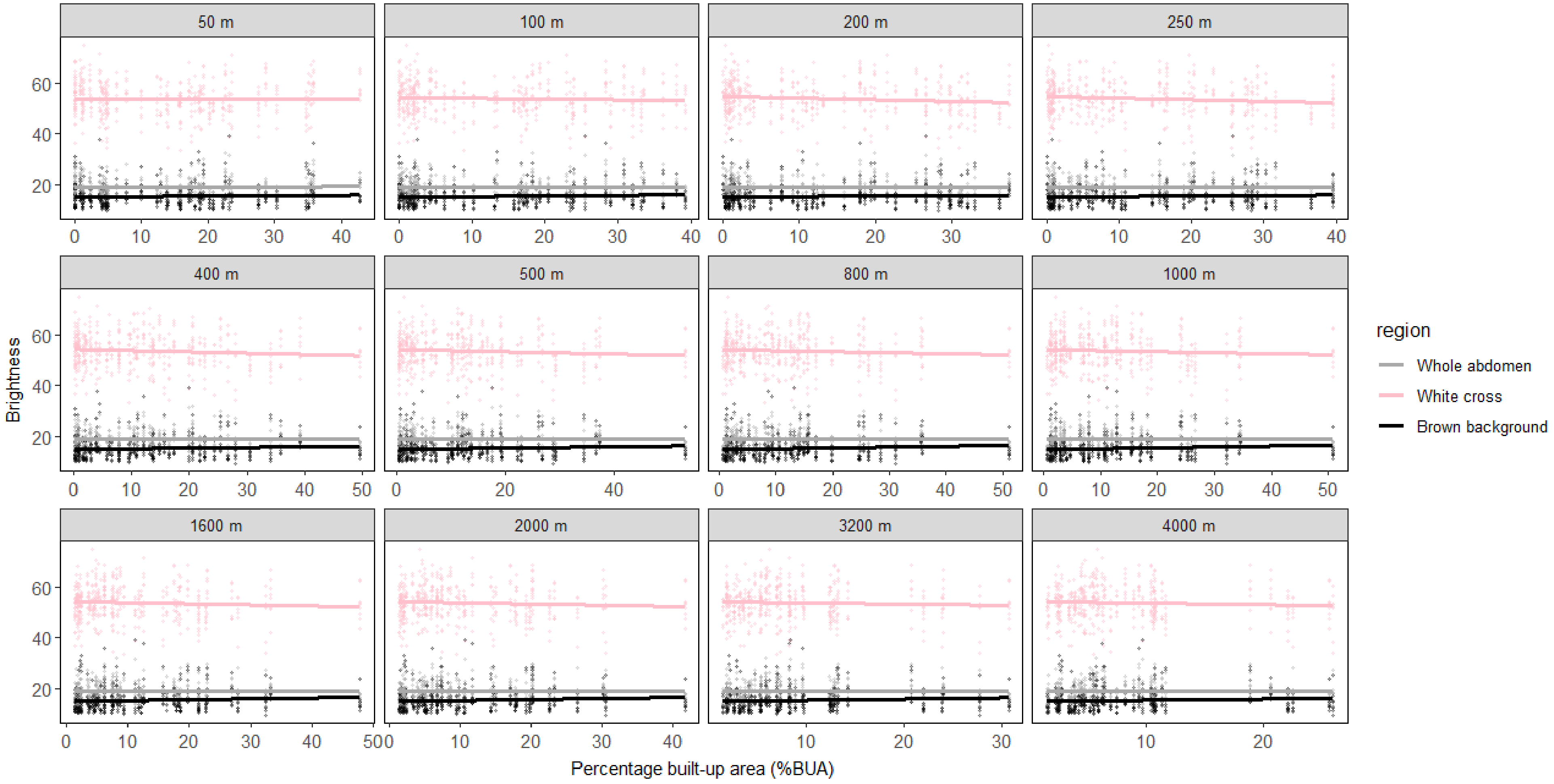


**Figure S5 Abdominal colour variation across the urbanization gradient.**

Brightness of the three abdominal regions (whole abdomen, white cross and brown background) plotted across the urbanization gradient (%BUA). Points represent individual measurements (n = 455). The white cross was not visible for three individuals (n = 452).

### Supporting Tables

**Table S1. Output of the linear mixed regression models and the model selection.**

| Variable | | *Spatial scale (radii in metres)* | | | | | | | | | | | |
|---|---|---|---|---|---|---|---|---|---|---|---|---|---|
| | | *50* | *100* | *200* | *250* | *400* | *500* | *800* | *1000* | *1600* | *2000* | *3200* | *4000* |
| Body length | %BUA | -0.064 (0.086) | 0.003 (0.086) | 0.098 (0.085) | 0.151 (0.083) | **0.208 (0.081)** | **0.214 (0.08)** | **0.229 (0.08)** | **0.252 (0.079)** | **0.283 (0.076)** | **0.284 (0.076)** | **0.317 (0.074)** | **0.339 (0.072)** |
| | AICc | 1214.5 | 1215.1 | 1213.8 | 1211.9 | 1208.9 | 1208.4 | 1207.4 | 1205.7 | 1202.9 | 1202.8 | 1199.2 | **1196.6** |
| Abdomen area | %BUA | -0.127 (0.088) | -0.065 (0.089) | 0.033 (0.09) | 0.087 (0.089) | 0.153 (0.087) | 0.159 (0.087) | **0.174 (0.087)** | **0.195 (0.086)** | **0.234 (0.084)** | **0.236 (0.084)** | **0.279 (0.081)** | **0.309 (0.079)** |
| | AICc | 1192.4 | 1193.9 | 1194.3 | 1193.5 | 1191.4 | 1191.2 | 1190.6 | 1189.5 | 1187.2 | 1187.0 | 1183.7 | **1181.0** |
| Abdomen area (log-transformed; body length as covariate)[a] | %BUA | **-0.056 (0.025)** | **-0.071 (0.025)** | **-0.071 (0.025)** | **-0.069 (0.025)** | **-0.061 (0.025)** | **-0.059 (0.025)** | **-0.062 (0.025)** | **-0.063 (0.025)** | **-0.058 (0.025)** | **-0.058 (0.025)** | **-0.052 (0.026)** | -0.045 (0.026) |
| | AICc | 214.5 | **211.5** | 211.6 | 211.9 | 213.7 | 214.1 | 213.6 | 213.3 | 214.3 | 214.3 | 215.2 | 216.3 |
| Cross prominence index | %BUA | -0.012 (0.053) | -0.015 (0.053) | -0.014 (0.053) | -0.028 (0.053) | -0.063 (0.052) | -0.06 (0.052) | -0.054 (0.052) | -0.062 (0.052) | -0.071 (0.052) | -0.068 (0.052) | -0.078 (0.052) | -0.084 (0.052) |
| | AICc | 1226.3 | 1226.3 | 1226.3 | 1226.1 | 1224.9 | 1225.1 | 1225.3 | 1225.0 | 1224.5 | 1224.7 | 1224.1 | **1223.8** |
| Abdominal brightness* | %BUA | 0.026 (0.088) | 0.011 (0.087) | -0.026 (0.088) | -0.025 (0.088) | -0.01 (0.087) | -0.01 (0.088) | 0.002 (0.087) | 0.005 (0.088) | -0.006 (0.087) | -0.007 (0.087) | -0.009 (0.087) | -0.018 (0.087) |
| | AICc | 1206.8 | 1206.9 | **1206.8** | 1206.8 | 1206.9 | 1206.9 | 1206.9 | 1206.9 | 1206.9 | 1206.9 | 1206.9 | 1206.9 |
| White cross brightness* | %BUA | -0.011 (0.073) | -0.061 (0.073) | -0.12 (0.071) | -0.118 (0.071) | -0.094 (0.072) | -0.083 (0.072) | -0.071 (0.072) | -0.068 (0.073) | -0.07 (0.072) | -0.07 (0.072) | -0.055 (0.073) | -0.064 (0.073) |
| | AICc | 1253.0 | 1252.4 | **1250.3** | 1250.4 | 1251.4 | 1251.7 | 1252.1 | 1252.2 | 1252.1 | 1252.1 | 1252.5 | 1252.3 |
| Brown background brightness* | %BUA | 0.034 (0.079) | 0.06 (0.079) | 0.049 (0.079) | 0.052 (0.079) | 0.069 (0.078) | 0.067 (0.079) | 0.081 (0.078) | 0.089 (0.078) | 0.084 (0.078) | 0.087 (0.078) | 0.083 (0.078) | 0.069 (0.078) |
| | AICc | 1243.8 | 1243.4 | 1243.6 | 1243.5 | 1243.2 | 1243.2 | 1242.9 | **1242.7** | 1242.8 | 1242.7 | 1242.8 | 1243.2 |
| $\Delta T_{webhub-ambient}$ | %BUA | 0.049 (0.055) | 0.052 (0.055) | 0.038 (0.055) | 0.026 (0.056) | 0.034 (0.055) | 0.029 (0.055) | 0.025 (0.055) | 0.017 (0.056) | 0.001 (0.055) | 0.001 (0.056) | -0.014 (0.055) | -0.021 (0.056) |
| | AICc | 1293.4 | **1293.3** | 1293.8 | 1294.0 | 1293.8 | 1294.0 | 1294.0 | 1294.1 | 1294.2 | 1294.2 | 1294.2 | 1294.1 |
| $\Delta T_{retreat-ambient}$ | %BUA | 0.059 (0.059) | 0.079 (0.058) | 0.086 (0.058) | 0.077 (0.058) | 0.068 (0.059) | 0.065 (0.059) | 0.047 (0.059) | 0.033 (0.059) | 0.021 (0.059) | 0.018 (0.059) | 0.012 (0.059) | 0.018 (0.059) |
| | AICc | 1288.8 | 1287.9 | **1287.6** | 1288.1 | 1288.4 | 1288.6 | 1289.1 | 1289.4 | 1289.6 | 1289.7 | 1289.7 | 1289.7 |
| $\Delta T_{spider-webhub}$ | %BUA | 0.011 (0.093) | -0.028 (0.092) | 0.01 (0.092) | 0.03 (0.092) | 0.079 (0.091) | 0.096 (0.091) | 0.104 (0.09) | 0.11 (0.09) | 0.108 (0.09) | 0.09 (0.09) | 0.042 (0.09) | 0.026 (0.09) |
| | AICc | 763.1 | 763.0 | 763.1 | 763.0 | 762.4 | 762.0 | 761.8 | **761.7** | 761.7 | 762.1 | 762.9 | 763.0 |

| | | | | | | | | | | | | | |
|---|---|---|---|---|---|---|---|---|---|---|---|---|---|
| $\Delta T_{spider-retreat}$ | %BUA | -0.152 (0.091) | -0.206 (0.09) | -0.207 (0.091) | -0.204 (0.091) | -0.168 (0.092) | -0.152 (0.092) | -0.103 (0.094) | -0.083 (0.095) | -0.04 (0.096) | -0.027 (0.096) | -0.035 (0.097) | -0.043 (0.098) |
| | AICc | 485.0 | **482.8** | 482.8 | 483.0 | 484.5 | 485.1 | 486.5 | 487.0 | 487.6 | 487.7 | 487.6 | 487.5 |
| Thermal offset[b] | %BUA | 0.003 (0.076) | -0.043 (0.075) | --0.037 (0.076) | -0.032 (0.076) | 0.001 (0.076) | 0.01 (0.076) | 0.022 (0.075) | 0.027 (0.075) | 0.034 (0.075) | 0.030 (0.075) | 0.002 (0.075) | -0.007 (0.076) |
| | AIC | 1246.8 | **1246.5** | 1246.6 | 1246.6 | 1246.8 | 1246.8 | 1246.7 | 1246.7 | 1246.6 | 1246.7 | 1246.8 | 1246.8 |

AIC-summary statistics of the model selection and effect estimates (estimate and standard error) for all models testing the relationship between %BUA (calculated at radii of 50-4000 m) and microclimate selection, behavioural thermoregulation, thermal offset, body size and body colouration. Models with $\Delta AIC < 2$ are indicated as they support the data equally good as the model with the lowest AIC . Models with significant effects ($p < 0.05$) of %BUA are shown in bold.

*brightness was log transformed.

a: body size–corrected abdomen area was modelled as log-transformed abdomen area and log-transformed body length was included as a covariate.

b: position spider in the web (web hub or retreat) included as a covariate.

**Table S2: Posterior estimates from the Bayesian multivariate mixed models jointly modelling body size and colouration.** Body size (body length) and abdominal brightness were modelled jointly as a function of %BUA measured at radii from 50 to 4000 m. A null model without urbanization was also fitted for comparison. The estimated effect of urbanization on body size increased with spatial scale, consistent with the univariate analyses, whereas abdominal brightness showed no detectable response. Estimated among-site (random-effect) and within-site (residual) correlations between body size and abdominal brightness remained highly consistent across scales (and in the null model), indicating that the inferred covariance structure was robust to the spatial scale at which urbanization was quantified. Model support was compared using leave-one-out cross-validation (LOO). Values are shown as differences in expected log predictive density (ΔELPD) relative to the best-supported model, together with the standard error of the difference (SE).

| radius (m) | Body size effect (β) | 95% CrI | Colour effect (β) | 95% CrI | Among-site (random-effect) correlation | 95% CrI | Within-site (residual) correlation | 95% CrI | Δ ELPD ± SE |
|---|---|---|---|---|---|---|---|---|---|
| None (no urbanization predictor) | / | / | / | / | -0.43 | 0.68, -0.13 | 0.22 | 0.13, 0.31 | -2.2 ± 3.1 |
| 50 | -0.06 | -0.24, 0.11 | 0.03 | -0.15, 0.21 | -0.43 | -0.69, -0.11 | 0.22 | 0.13, 0.31 | -2.7 ± 3.1 |
| 100 | 0.00 | -0.17, 0.18 | 0.01 | -0.17, 0.20 | -0.42 | -0.69, -0.12 | 0.22 | 0.13, 0.31 | -2.4 ± 3.0 |
| 200 | 0.09 | -0.07, 0.26 | -0.02 | -0.20, 0.16 | -0.44 | -0.69, -0.13 | 0.22 | 0.13, 0.31 | -3.2 ± 2.8 |
| 250 | 0.15 | -0.02, 0.31 | -0.03 | -0.21, 0.15 | -0.44 | -0.71, -0.13 | 0.22 | 0.13, 0.31 | -3.3 ± 2.6 |
| 400 | 0.20 | 0.04, 0.38 | -0.01 | -0.18, 0.18 | -0.46 | -0.72, -0.16 | 0.22 | 0.13, 0.31 | -2.6 ± 2.3 |
| 500 | 0.21 | 0.05, 0.38 | -0.01 | -0.19, 0.16 | -0.46 | -0.72, -0.15 | 0.22 | 0.13, 0.32 | -1.8 ± 2.2 |
| 800 | 0.23 | 0.07, 0.40 | 0.00 | -0.18, 0.17 | -0.48 | -0.74, -0.17 | 0.22 | 0.13, 0.31 | -1.4 ± 2.1 |
| 1000 | 0.25 | 0.10, 0.41 | 0.01 | -0.17, 0.19 | -0.49 | -0.74, -0.19 | 0.22 | 0.13, 0.31 | -1.7 ± 1.9 |
| 1600 | 0.28 | 0.13, 0.44 | -0.01 | -0.19, 0.18 | -0.50 | -0.75, -0.19 | 0.22 | 0.13, 0.31 | -0.7 ± 1.5 |
| 2000 | 0.28 | 0.13, 0.44 | -0.01 | -0.19, 0.17 | -0.50 | -0.75, -0.20 | 0.22 | 0.13, 0.31 | -1.5 ± 1.3 |
| 3200 | 0.32 | 0.16, 0.47 | -0.01 | -0.19, 0.17 | -0.52 | -0.77, -0.22 | 0.22 | 0.13, 0.31 | -0.1 ± 0.5 |
| **4000** | **0.34** | **0.19, 0.49** | **-0.01** | **-0.18, 0.16** | **-0.51** | **-0.78, -0.19** | **0.24** | **0.15, 0.33** | **0.0 ± 0.0** |

**Table S3. Individuals exhibiting the highest and lowest brightness values for whole-abdomen, white cross and brown background colouration.**

These identifiers are provided to document the range of colour variation present in the dataset.

| Part of the abdomen | Spider with highest brightness | Spider with lowest brightness |
|---|---|---|
| Whole abdomen | SC22P14SR09<br>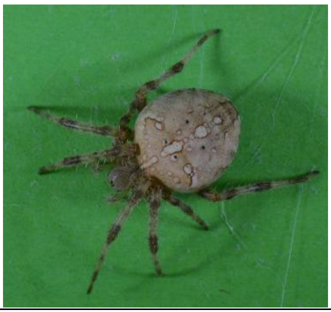 | SC22P13SR01<br>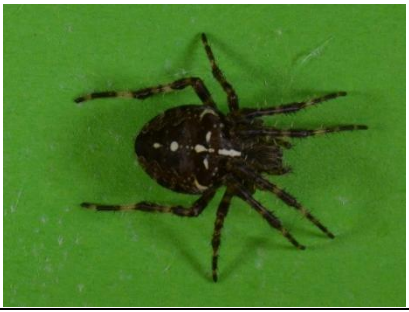 |
| White cross | SC22P18SG08<br>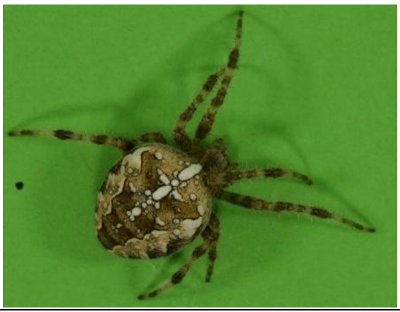 | SC22P21SG06<br>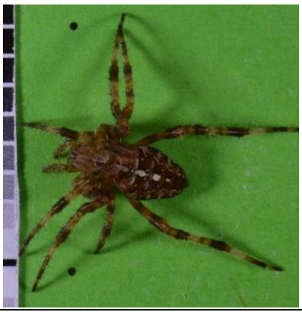 |
| Brown background | spider SC22P14SR09<br>Second brightest individual :<br>SC22P09SG04<br>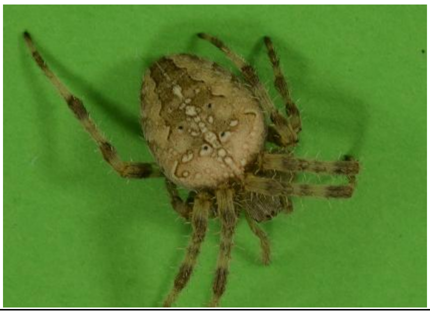 | SC22P20SR04<br>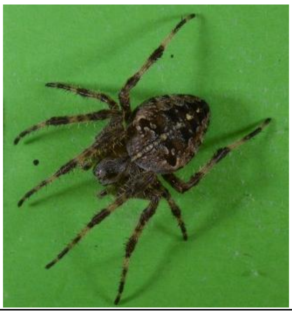 |
